# Synergistic effect of heat and drought on leaf VOC emissions and root exudates in Norway spruce saplings

**DOI:** 10.64898/2026.04.29.721567

**Authors:** Melissa Wannenmacher, Mirjam Meischner, Clara Stock, Stefanie Dumberger, Jürgen Kreuzwieser, Simon Haberstroh, Christiane Werner

**Affiliations:** Ecosystem Physiology, Faculty of Environment and Natural Resources, University of Freiburg, Georges-Köhler-Allee 53/54, 79110 Freiburg im Breisgau, Germany

**Keywords:** Root exudates, volatile organic compounds (VOC), *Picea abies*, compound drought, stress interaction, drought, heat

## Abstract

Compound droughts, i.e. the co-occurrences of heat and drought, represent a serious challenge for temperate forest trees leading to significant losses in forest biomass.

We studied the physiological response of Norway spruce (*Picea abies*) saplings to heat and drought individually, and in combination. Continuous measurements of leaf gas exchange and VOC emission allowed us to identify fast-response reactions, while discrete VOC and root exudate samplings added qualitative information on compositional changes. Additionally, we used ^13^CO_2_ and ^2^H_2_O label pulses to investigate C-allocation and root water uptake in response to stress.

Heat as well as drought reduced assimilation rates in the saplings, whereas transpiration, leaf VOC emission and root exudation rates increased in response to heat. Drought alone increased VOC emission but decreased exudation rates. Combined heat and drought triggered an amplified response in both processes despite negative net CO_2_ assimilation rates. Label incorporation showed compromised water uptake capacity of drought-stressed plants and illustrated *de novo* C-allocation to VOC emission and root exudates.

The results point at the high susceptibility of Norway spruce saplings to drought and heat. Combined stress resulted in synergistic responses in VOC emissions and root exudates, showing the detrimental effect of compound droughts on Norway spruce.

**Highlight:** In this study, we found synergistic effects of heat and drought on carbon losses from leaf VOC emission and root exudates despite negative assimilation rates in Norway spruce saplings.

## Introduction

Plants face a trade-off withstanding combined heat and drought: Increased or maintained transpiration under heat costs precious water, whereas closing stomata to avoid excessive water loss leads to rising, potentially critical leaf temperatures (Kunert *et al*., 2022). These compound droughts, i.e. the co-occurrences of heat and drought, are expected to increase in frequency and severity (Bevacqua *et al*., 2022) causing novel constrains to plant functioning (Werner *et al*., 2025). Economically relevant tree species of European temperate forests are not well adapted to these conditions. Hence, compound droughts have already caused 46% of tree mortality in Europe (Gazol and Camarero, 2022). Norway spruce (*Picea abies*) is one key species of European forests with significant economic value (Knoke *et al*., 2021). The species has already experienced significant losses in vitality and abundance due to climate change (Arend *et al*., 2021) and future declines are expected (Kolář *et al*., 2017). To predict future consequences of continuously changing environmental conditions, such as increased frequency and severity of compound droughts, a thorough understanding of plant-physiological processes in Norway spruce is crucial.

There are several strategies for trees to acclimate to heat and drought. One of them is the adjustment of leaf gas exchange via stomatal regulation (Eamus *et al*., 2008). Norway spruce has shown to act as rather isohydric species closing stomata early under increasingly drier condition and recovering only slowly from drought events (Ulrich and Grossiord, 2023; Hesse *et al*., 2024; Karpov *et al*., 2026). This strategy induces elevated leaf temperatures due to reduced transpiration cooling (Eamus *et al*., 2008), especially when drought is complemented by heat. The rather long recovery time of Norway spruce after drought (Ulrich and Grossiord, 2023) increases its risk in case of recurrent extreme events. Seedlings and saplings are particularly prone to drought stress due to lower root biomass and internal water storage compared to mature trees, which have a more balanced water status (Oberhuber *et al*., 2015).

The emission of volatile organic compounds (VOCs) represents another strategy to cope with heat and drought. Once released, VOCs can alter the atmospheric chemistry by increasing the lifetime of greenhouse gases and by favouring the formation of ozone and particulate matter (Atkinson and Arey, 2003; Rennenberg *et al*., 2006). Mature Norway spruce trees emit 0.05 - 0.5% of their photosynthetically fixed C in the form of VOCs (Grabmer *et al*., 2006). Under stress conditions, however, this can rise substantially, particularly when photosynthesis declines (Grabmer *et al*., 2006). Norway spruce VOC emissions are dominated by terpenoids, with varying shares of mono- and sesquiterpenoids depending on environmental conditions (Hakola *et al*., 2017; Duan *et al*., 2020; Meischner *et al*., 2024) and act as signalling molecules mediating plant-plant interaction (Holopainen, 2004; Meischner *et al*., 2025). They are known to increase the stress tolerance of plants (Peñuelas *et al*., 2005; Kreuzwieser *et al*., 2021; Meischner *et al*., 2024). It has been shown that VOC production allows quenching of excess energy or metabolites, which might occur under heat or drought (Loreto and Schnitzler, 2010; Werner *et al*., 2020). Furthermore, isoprene and monoterpenes are known to induce thermo-stabilization of membranes under elevated temperature (Sharkey *et al*., 2008) and to reduce the damage through reactive oxygen species (Loreto and Schnitzler, 2010). VOC emissions increase exponentially with increasing temperature (Rennenberg *et al*., 2006; Grabmer *et al*., 2006; Bourtsoukidis *et al*., 2024). This is partly due to an increased vapour pressure of VOCs under heat and especially relevant in VOC storing species like Norway spruce. In contrast, several studies have found decreased VOC emissions in Norway spruce under drought, which might be a result of resource limitation (Filella *et al*., 2007; Haberstroh *et al*., 2018; Daber *et al*., 2025). However, it has been suggested that the drought intensity plays a crucial role, where emissions increase upon the onset of drought and decline under prolonged or more severe drought (Ormeño *et al*., 2007; Wu *et al*., 2015; Bonn *et al*., 2019), or the composition of VOCs shifts (Werner *et al*., 2021). Combined stressors or prolonged exposure to one stressor can lead to reduced VOC emissions due to a declining net CO_2_ assimilation (Ormeño *et al*., 2007; Meischner *et al*., 2024). Yet, the effect of combined drought and heat on VOC emissions in Norway spruce has not been studied intensively to date.

Analogous to aboveground VOC emissions, root exudates constitute a belowground carbon loss for plants acclimating to environmental stress. Root exudates are known to facilitate soil penetration and nutrient uptake and can therefore mitigate effects of drought (Gargallo-Garriga *et al*., 2018). However, research on the effect of drought on root exudates led to ambiguous results (Heinzle *et al*., 2023). While some studies found enhanced exudation under drought (Leuschner *et al*., 2022; Brunn *et al*., 2022), others observed a reduction (Dannenmann *et al*., 2009; Li *et al*., 2025). Possible reasons for these contradictory findings could be species-specific responses to environmental conditions (Wannenmacher *et al*., 2025) or differences in extend of drought between studies. Drought intensity and duration might be decisive as suggested by Gargallo-Garriga *et al*. (2018), indicating that exudation is increased in an initial state of drought and decreases thereafter, a similar pattern as observed for VOC emission. Data in higher temporal resolution is therefore needed to elucidate exudation dynamics under drought. Regarding the effect of temperature on root exudates, there is larger consensus among several studies showing increased exudation rates under elevated temperatures (Rohrbacher and St-Arnaud, 2016; Meier *et al*., 2020; Leuschner *et al*., 2022; Heinzle *et al*., 2023). This might be due to a higher share of non-structural carbohydrates in the roots due to increased photosynthetic activity under warmer conditions (Prescott *et al*., 2020). However, recent research suggests that increased exudation rates are a short-term response of plants to heat, while under prolonged periods of higher temperature, exudation rates remain unaltered (Heinzle *et al*., 2023; Liu *et al*., 2025). Thus, there is still a large uncertainty regarding the dynamics in exudation patterns, the understanding of which is crucial to predict the resilience of plants and the fate of C under a warming climate.

The objectives of this study are to (1) investigate the temporal dynamics of leaf VOC emission and root exudates in Norway spruce saplings under heat, drought and their interaction. We aim at (2) fostering the understanding of how Norway spruce prioritises above and belowground processes when exposed to single or combined stress. We continuously measured leaf gas exchange and used stable isotope ^13^CO_2_ and ^2^H_2_O labelling to (3) investigate water uptake dynamics and allocation of *de novo* synthesised C to VOC emission and root exudates. We hypothesise that C-allocation into VOC emission and root exudates increases under initial exposure to a single stressor, but decreases under prolonged or combined stress, when C becomes limiting. Further, we expect that exudation from roots is favoured under drought to compensate for compromised root-soil contact, whereas VOC emissions are favoured under heat due to their positive effect on heat tolerance.

## Material and Methods

### Plant material

Eight five-year old Norway spruce saplings originating from a nursery in Southeast Germany (Baumschulen Haage GmbH & Co. KG in Langerringen, Bavaria, Germany) were planted into 30.5 l boxes in spring 2023, more than one year before the experiment started. To avoid waterlogging, boxes were equipped with small holes for drainage and a layer of expanded clay aggregate at the bottom (∼4.5 l). On top of the expanded clay aggregate, 24 l of fertilised potting soil (Breisgau Kompost, Müllheim, Germany) mixed with sand (grain size ∼1 mm; Holcim Kies und Beton GmbH, Hamburg, Germany) were added (5:1, soil:sand). The boxes had one open side, which was covered by a removable acrylic glass pane (Ketterer + Liebherr GmbH, Freiburg, Germany) so that roots could be accessed for exudate sampling. To avoid light disturbance of the roots, glass panes were covered by a 0.5 mm dark tarpaulin. Plants were watered manually throughout the experiment.

### Experimental design

Half of the saplings were exposed to drought stress (drought-stressed plants, n = 4), while the other half was supplied with sufficient water (control plants, n = 4). To ensure adequate soil moisture conditions for the two groups from the beginning of the heat stress experiment, water supply to drought-stressed plants was stopped two months before the start of the heat stress experiment to mimic the drought development under natural conditions (Fig. 1). All plants were kept outside under semi-controlled conditions under a 100% rainout shelter for the eight weeks of drought treatment, while the ten-day heat stress experiment took place under fully controlled conditions in a walk-in climate chamber (ThermoTEC, Weilburg, Germany). Leaf gas exchange, leaf water potential (Ψ_leaf_) and root exudate measurements were conducted once before and regularly during the drought treatment in June and July to monitor the plant status. Saplings were moved to the climate chamber one week before the start of the heat stress experiment for acclimatisation. In the climate chamber, environmental parameters were fully controlled (temperature: 25°C during daytime, 15°C during night-time; photoperiod: 15.5 h light, 7.5 h dark, dawn and dusk: 30 min each; light intensity: 600 µmol m^-2^ s^-1^; relative humidity: 60%) and plants were watered every two to three days (200 ml for drought-stressed plants; sufficient to control plants). Continuous measurements of leaf gas exchange, including isotopic signatures of δ^2^H_2_O and δ^13^CO_2_, and VOC emission rates in the climate chamber started on 3 August 2024, referred to as day 0 in the climate chamber (see Fig. 1). On day 3 in the climate chamber, the air temperature was increased from 25 to 35°C by steps of 2°C every two hours (heat ramp) and the heat (35°C during daytime, 25°C during night-time) was kept for nine days (i.e., until day 12 in the climate chamber). The plants were supplied with ∼20 ppm of 99% ^13^CO_2_ added to the cuvette inflow for 4 h on day 5 in the climate chamber, resulting in a label strength of ∼353‰ δ^13^CO_2_. One day later, all plants were supplied with 200 ml of water enriched in ^2^H (δ^2^H_2_O: 5295‰ ± 64‰). To determine emitted terpenoids and their carbon isotopic composition, four additional VOC emission measurements for subsequent GC-MS-IRMS analysis took place during the heat stress experiment. Exudation measurements were performed four days and eight days after the heat ramp. Ψ_leaf_ was measured on day 12 in the climate chamber at the end of the experiment.

**Figure 1:**
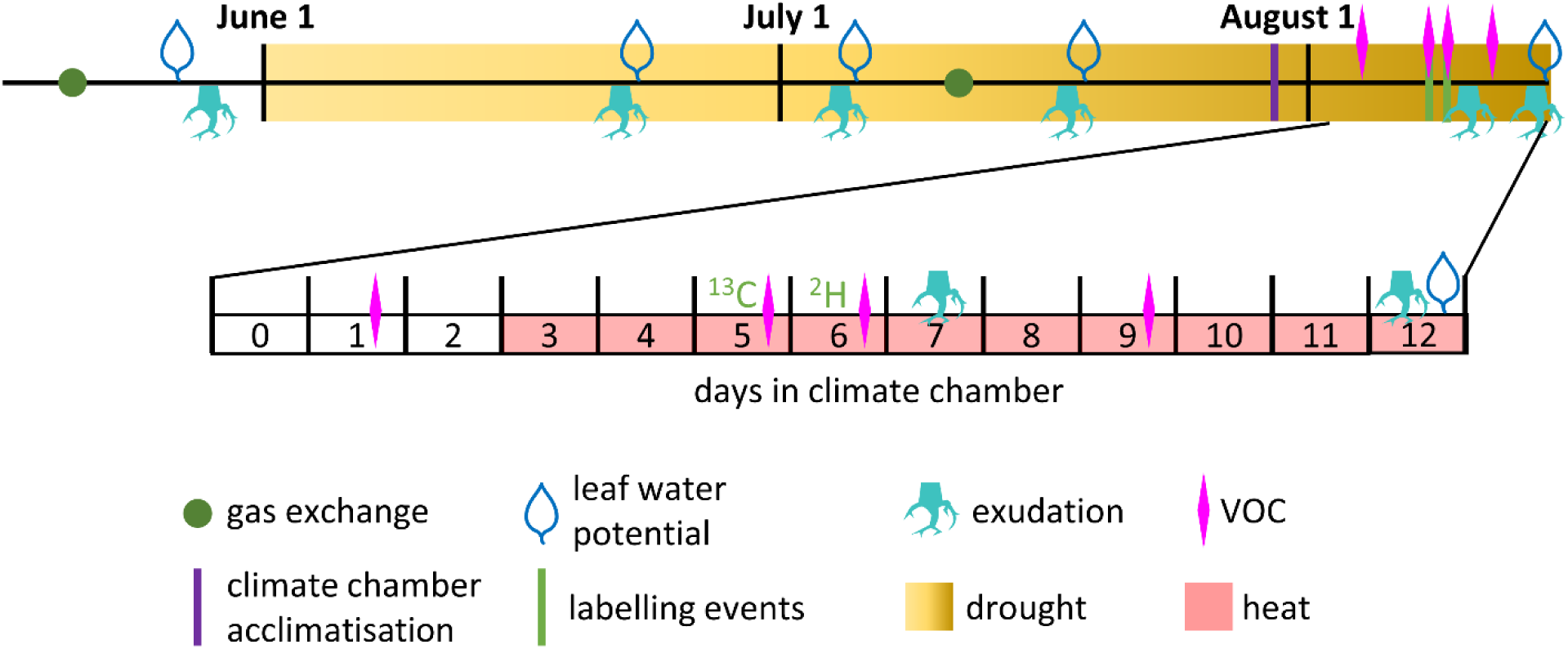
Timeline illustrating samplings, measurements, labelling events and the most important changes in growing conditions during the drought treatment and the heat stress experiment. Note that control plants were not exposed to drought.

### Meteorological data

During the drought treatment, air temperature was measured (S-THC-M002, Onset Computer Corporation, Bourne, USA) ∼1 m above the canopy under the rainout shelter with a measurement interval of 5 min and recorded by a HOBO Micro Station Data Logger (Onset Computer Corporation, Bourne, USA). Soil volumetric water content (VWC) in every box (n=8) was recorded in 5 min intervals by soil moisture sensors 5TM and GS1 connected to data loggers EM50 (all Decagon Devices Inc., Pullman, USA). The relative soil volumetric water content (VWC_rel_) at time point *i* was calculated based on minimum VWC (*VWC_min_*) and maximum VWC (*VWC_max_*) recorded throughout the experiment according to the following formula:

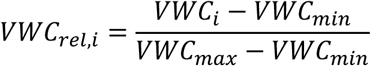

### Ecophysiological measurements during the drought treatment

Leaf water potential (Ψ_leaf_) was determined using a Scholander pressure chamber (Model 3000 Plant Water Status Console, Soil Moisture Equipment Corp., Santa Barbara, USA). Measurements were conducted around midday (between 13.00 and 14.00 MESZ) on sun-lit needles once before the drought treatment, every two weeks during the drought treatment and once on day 12 in the climate chamber.

Leaf gas exchange during the drought treatment was measured using a portable gas exchange measuring system (GFS 3000; Heinz Walz GmbH, Effeltrich, Germany) with an air temperature of 23°C, a relative humidity of 35% and a light intensity of 600 µmol m^-2^ s^-1^. The CO_2_ concentration in the measuring cuvette was adjusted to 430 ppm. Measurements were conducted once before and once under moderate drought intensity around midday between 10.30 and 15.30 MESZ.

### Continuous gas exchange and VOC emission measurement

After the drought treatment, plants were moved to the climate chamber and acclimated for one week. Continuous VOC fluxes and isotope discriminated gas exchange was measured in the climate chambers with our automatic cuvette switching unit (Fasbender *et al*., 2018; Werner *et al*., 2020). In short, branches of the eight Norway spruce saplings were enclosed in air-tight cuvettes consisting of Nalophan foil and equipped with a separate inlet and outlet. In addition, there were two empty cuvettes for background measurements. All cuvettes were equipped with a fan to ensure an equilibrated gas mixture in the cuvette. Air entering the cuvettes was channelled through a zero-air generator and a CO_2_ absorber, then ∼430 ppm CO_2_ (δ^13^CO_2_ = -35.26 ± 0.03‰) were adjusted. The air from the cuvette outlet was directed to the analyser unit, consisting of (i) an isotope laser spectrometer (DeltaRay, Thermo Fisher Scientific Inc., Waltham, USA) to measure CO_2_ concentration and δ^13^CO_2_ isotopic signature, (ii) a water isotope laser spectrometer (L2130-i, Picarro Inc., Santa Clara, USA) to measure the water vapour content and δ^2^H_2_O isotopic signature, (iii) an infrared gas analyser (LI850, LI-COR Biotechnology, Lincoln, USA) to measure CO_2_ and H_2_O in the outlet air and (iv) a proton-transfer-reaction time-of-flight mass spectrometer (PTR-TOF-MS 4000; Ionicon Analytik Ges.m.b.H., Innsbruck, Austria) to determine VOC fluxes. The PTR-TOF-MS was operated with a drift tube temperature of 80°C, a drift voltage of 500 V and a drift pressure of 2.7 mbar (Meischner *et al*., 2024). The cuvettes were connected to a multi-position valve, so that the eight branch and two blank cuvettes were measured alternatingly for 7 min each. Net CO_2_ assimilation and transpiration rates were calculated according to Caemmerer and Farquhar (1981). Isotopic signatures δ^13^CO_2_ in respiration and δ^2^H_2_O in transpiration were calculated using the following equation:

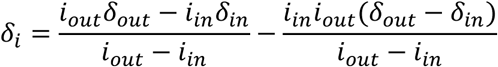

where *i* is the mole fraction of CO_2_/H_2_O and *δ* the isotopic ratio δ^13^CO_2_/δ^2^H_2_O of cuvette inflow (*in*) and outflow (*out*) (Dubbert *et al*., 2014). δ^2^H_2_O values were normalised by the mean isotopic signature in daytime transpiration of control plants during the three days before labelling, and expressed as Δδ^2^H_2_O, i.e. the difference of pre-label and post-label values.

### Continuous PTR-TOF-MS VOC data analysis

With the PTR-TOF-MS, VOC emissions were analysed in real-time based on their molecular mass (Jordan *et al*., 2009). The GLOVOCS-database (Yáñez-Serrano *et al*., 2021) was used to assign the measured protonated mass-to-charge ratios to individual compounds. Compounds of the same mass, such as different monoterpenes, were assigned to the same compound group. For quantification, a multi-component gas mixture (Apel Riemer Environmental, USA) in combination with a liquid calibration unit (Ionicon Analytic, Innsbruck, Austria) was used. The raw data was processed with the IDA software (version 239 2.2.0.7, Ionicon, Innsbruck, Austria). For each 7-min measurement interval per cuvette, mean emission rates were calculated, whereby the first 2 min were excluded to guarantee stable measurement conditions, and blank measurements were subtracted from plant measurements. All VOC emission rates were normalised by the cuvette flow rate through and based on projected leaf area.

### Compound-specific discrete terpenoid sampling and analysis

VOC terpenoid sampling took place four times during the experiment on glass thermodesorption tubes filled with Tenax TA connected to the outlets of the cuvettes and an air sampling pump (Pocket Pump TOUCH, SKC Inc., Pittsburgh, USA) at 120 ml min^-1^ for 1 h. Sampling took place in the afternoon of day 1 (pre-heat), day 5 (during ^13^CO_2_ labelling; 2^nd^ day of heat), day 6 (3^rd^ day of heat) and day 9 in the climate chamber (6^th^ day of heat). Subsequent VOC analysis was conducted by gas chromatography-mass spectrometry (GC-MS 7890B GC System & 5975C MSD, Agilent Technology, Santa Clara, USA) coupled to a combustion interface and an isotope ratio mass spectrometer (C-IRMS, GC5 & Isoprime PrecisION, Elementar Analysesysteme GmbH, Langenselbold, Germany) according to Haberstroh *et al*. (2019) and Kreuzwieser *et al*. (2021). Compound identification and peak area quantification was performed using the MASSHUNTER software (Agilent Technologies, Santa Clara, USA). Compounds were identified with the NIST library (National Institute of Standards and Technology) based on retention time and fragmentation patterns. Only compounds with a match factor > 90 were included in the analysis. Peak area integration by the software was checked and manually corrected when required. Peak areas detected in blanks were subtracted from plant samples. The following authentic standards were used for quantification: α-pinene, β-pinene, limonene, trans-β-ocimene and caryophyllene (all Sigma Aldrich, Merck group, Bulrington, USA). For monoterpenoids without authentic standard, a mean quantification factor was calculated from α-pinene, β-pinene, limonene, trans-β-ocimene. For sesquiterpenoids, the quantification factor of caryophyllene was used. A list of detected compounds, their compound groups and quantification standards can be found in the supplement (Table S1). Isotopic ratios from the IRMS were processed using the LyticOS software (Elementar Analysesysteme GmbH, Langenselbold, Germany).

### Root exudate sampling and analysis

The root exudate collection was based on the *in-situ* cuvette-based method described by Phillips *et al*. (2008) and conducted as in Wannenmacher *et al*. (2025). Roots were accessed through the removable glass panes and soil particles were carefully removed with water and tweezers. To recover from the cleaning and excavation process, roots were then wrapped into aluminium foil and left there for 24 hours. The aluminium foil was covered with soil to avoid light disturbance to the roots. Following the recovery time, the root was placed in a cuvette consisting of a 30 ml plastic syringe without plunger with Luer-Lock outlet (B.Braun, Melsungen, Germany) fitted with a 3-way-valve (Teqler, Wecker, Luxembourg). The outlet was covered with glass wool, and the cuvette filled with 3 mm glass beads to imitate the soil environment. 10 ml of a C-free dilute nutrient solution (0.5 mM NH_4_NO_3_, 0.1 mM KH_2_PO_4_, 0.2 mM K_2_SO_4_, 0.15 mM MgSO_4_*7H_2_O, 0.4 mM CaCl_2_*2H_2_O) were added. To avoid contamination, the cuvette was sealed with parafilm. 24 hours later, the nutrient solution was replaced by 5 ml of root exudate capturing solution, a C- and N-free dilute nutrient solution (0.1 mM KH_2_PO_4_, 0.2 mM K_2_SO_4_, 0.15 mM MgSO_4_*7H_2_O, 0.4 mM CaCl_2_*2H_2_O). Two flushes were performed before adding the capturing solution. 24 hours later, the capturing solution was retrieved and two flushes of 10 ml were performed to capture the exudates attached to the glass beads and the root. Retrieved solutions were immediately filtered through a 0.2 µm sterile syringe filter and stored at -80°C until analysis. The same procedure was applied to control cuvettes, which did not contain a root and later served as correction for contamination. The roots in the cuvettes were cut and cooled at ∼8°C until scanning. The root surface area was determined using the software WinRHIZO 2021a 32-Bit (Copyright 1993-2021, REGENT INSTRUMENTS INC) with an Epson Perfection V850 Pro Scanner (Seiko Epson Corporation, Tokyo, Japan). Samples were taken once before the drought treatment, every two weeks during the drought treatment and on days 7 and 12 in the climate chamber.

Frozen root exudate samples were freeze-dried for at least 80 h and then derivatised according to Maurer *et al*. (2021). In brief, the freeze-dried samples were solved in 100 µl of a 20 mg ml^-1^ methoxyamine hydrochloride in anhydrous pyridine solution. After centrifugation, 20 µl of the prepared solution were incubated on a thermoshaker at 30°C and 1400 rpm for 90 min. In a next step, 35 µl of N-methyl-N-(trimethylsilyl)-trifluoroacetamide (MSTFA) were added and samples went through shaking at 1400 rpm at 37°C for 30 min. Gas chromatography-mass spectrometry (see above) was used to analyse the derivatives with a split of 2:1. A detailed parameter description can be found in Kreuzwieser *et al*. (2009). The MASSHUNTER Quantitative Analysis software (Agilent Technologies, Santa Clara, USA) was used for peak detection, identification and peak area determination. Peak identification was based on the Golm Metabolome Database (2011) and peak area integration was, if necessary, corrected manually. Only compounds with a match factor larger than 50 were included in the analysis. To account for changes in sensitivity during the GC-MS measurement process, peaks were normalised by the internal standard ribitol. The following authentic standards were used for quantification: aspartic acid, boric acid, citric acid, cysteine, fructose, galactose, glucose, glutamic acid, glycerol, lactic acid, malic acid, N,N-dimethyl-glycine, phenylalanine, propane-1,3-diol, pyruvic acid, sucrose, thriethanolamine and uric acid (all Sigma Aldrich, Merck group, Bulrington, USA). The detected compounds were categorised in 10 groups: organic acids, amino acids, other acids, aromatic compounds with and without N, other N-containing compounds, alkanes, alcohols, sugars and unknown compounds. A detailed list of detected compounds, grouping and the standards used for quantification can be found in the supplement (Table S2). To determine the carbon isotopic signature δ^13^C in the root exudates, the freeze-dried samples were transferred into tin capsules and analysed by an Isoprime PrecisION isotope ratio mass-spectrometer (IRMS; Elementar Analysesysteme GmbH, Langenselbold, Germany; (Werner *et al*., 2009). A trap current of 200 µA CO_2_ (dilution 5%) was used. Tyrosine and caffeine IAEA-600 were used as standards with δ^13^C reference values of -24.80‰ for tyrosine and -27.77‰ for IEAE-600.

### Statistical analysis

Net CO_2_ assimilation, transpiration, leaf water potential and the isotopic signature of respiration, transpiration, and root exudates were evaluated using two-way analysis of variance to compare treatments and measurement dates. If needed, post-hoc comparisons were based on least-square means combined with a sidak-correction to find significant differences between treatments and measurement days. A significance level of α=0.05 was chosen for all analyses. Normal distribution of residuals and homogeneity of variance were tested with Shapiro-Wilk and Levene’s test. A locally estimated scatterplot smoothing (LOESS) regression was used in continuous net CO_2_ assimilation, transpiration and PTR-TOF-MS VOC measurement graphs.

Non-normally distributed VOC emission rates and isotopic signature as well as exudation rates were compared between treatments using the Kruskal-Wallis test and between measurement days with the Friedman test combined with the Conover Squared Rank test for post hoc analysis if needed.

All data were processed using R (version 4.5.1, R Core Team, 2025). Graphs were generated using the package ‘ggplot2’ (Wickham, 2016).

## Results

### Ecophysiological response of Norway spruce to soil drying

During the drought treatment under the rainout shelter, ambient air temperature varied between 8 and 35°C (Fig. 2 a). While air temperatures did not rise above 30°C until mid-June, values above 30°C were more common thereafter. The two days before the second exudation sampling were marked by a first sudden natural heat spell with a maximum temperature of 31.6°C. While the regular irrigation of control plants kept the VWC_rel_ at an intermediate level throughout the experiment, the VWC_rel_ of drought-stressed plants deceased continuously until the start of the heat stress experiment reaching a minimum mean of 0.10 ± 0.05. Watering of drought-stressed plants during the heat stress experiment slightly increased the VWC_rel_ and ensured the survival of the plants, but stayed well below the VWC_rel_ of control plants (Fig. 2 b).

**Figure 2:**
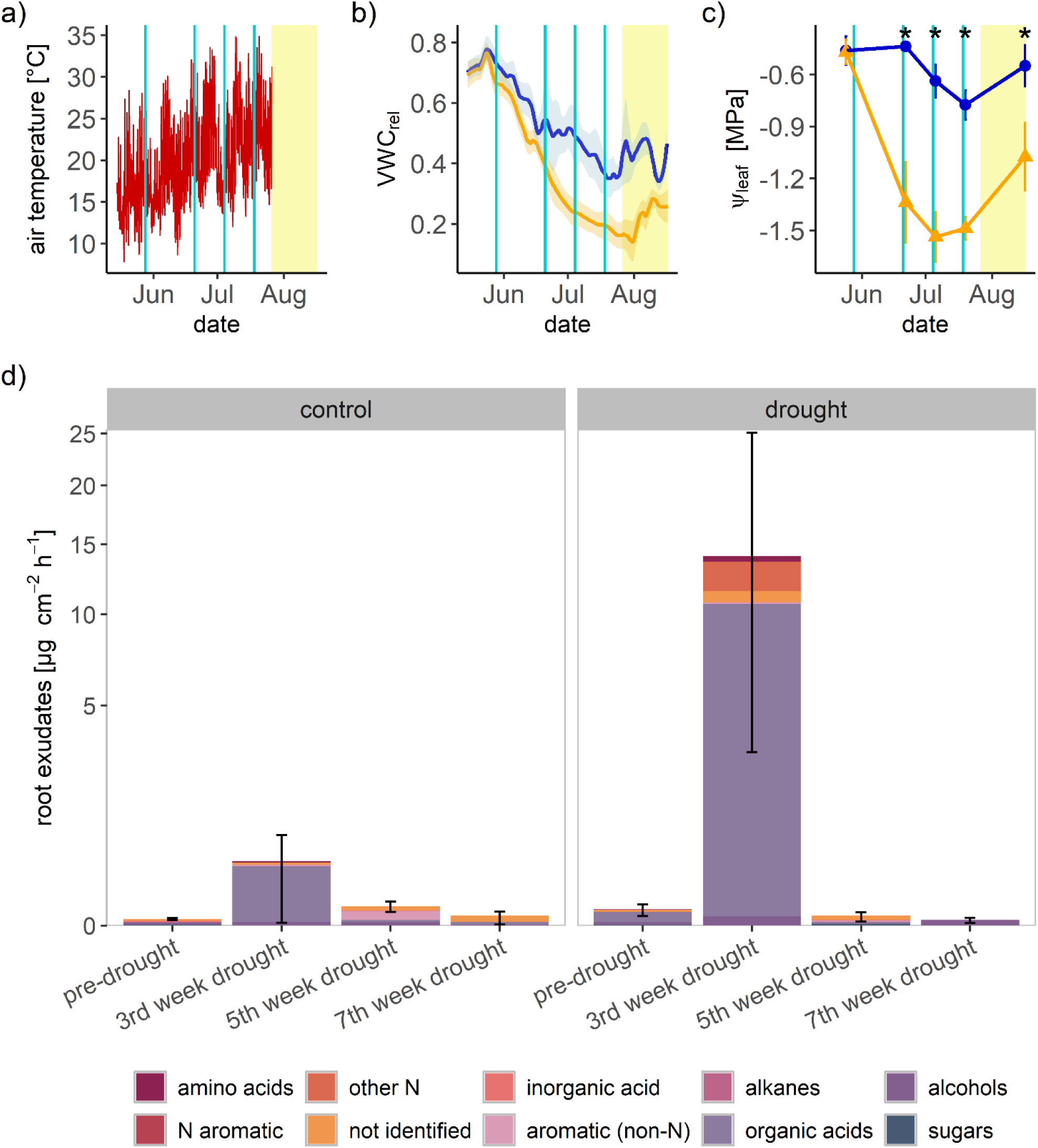
Air temperature (a), relative soil volumetric water content (VWC_rel_) (b), midday leaf water potential (Ψ_leaf_) for control (blue) and drought-stressed (orange) plants (c) and exudation rates from roots during the drought treatment (d). Areas shaded in yellow indicate the period in the climate chamber after the drought treatment. Turquoise lines in a – c indicate the time points of exudation samplings illustrated in d. Please note the square root scales on the y-axis for root exudates.

Net CO_2_ assimilation (*A*) and transpiration (*E*) rates were significantly reduced in drought-stressed plants in the fifth week of drought (*A* = 1.4 ± 0.2 µmol m^-2^ s^-1^, *E* = 0.2 ± 0.03 mmol m^-2^ s^-1^) compared to control plants (*A* = 3.9 ± 0.4 µmol m^-2^ s^-1^, *E* = 0.9 ± 0.1 mmol m^-2^ s^-1^) and pre-drought values in both treatments (Fig. S1). Before the drought treatment, all plants showed similar Ψ_leaf_ (-0.5 ± 0.2 MPa), but Ψ_leaf_ of drought-stressed plants were significantly lower (p = 0.03) already in the second week of drought, compared to control plants (Fig. 2 c), a difference that was maintained thereafter (-0.6 ± 0.2 MPa for control, -1.4 ± 0.4 MPa for drought-stressed saplings). A slight non-significant increase in Ψ_leaf_ in drought-stressed plants on day 12 in the climate chamber (-1.1 ± 0.2 MPa) was due to water supply to avoid plant mortality (Fig. 2 c).

A strong increase in exudation rates from roots from pre-drought to the third week of drought could be observed in both treatments (Fig. 2 d), with exudation rates increasing by a factor of ∼100 in control and a factor of >500 in drought-stressed plants. Peak exudation coincided with the first natural heat event of the season (Fig. 2 a). Here, organic acids dominated exudation composition with shares of 86% in control and 76% in drought-stressed plants. Moreover, there was a relatively high exudation rate of aromatic compounds compared to all other sampling dates. However, this effect was not significant. In the fifth and seventh week of drought, exudation rates were reduced again with minimum rates of 10 and 3 ng cm^-2^ h^-1^ in the seventh week of drought in control and drought-stressed plants, respectively.

### Climate chamber heat experiment: leaf gas exchange and VOC fluxes

Plants were transferred to the climate chamber and acclimated 10 days prior to the heat treatment. Rates of net CO_2_ assimilation (*A*) and transpiration (*E*) differed significantly between well-watered and drought-stressed plants in the pre-heat phase (Fig. 3 a and b). Heat led to an immediate drop in net assimilation in both treatments leading to negative mean daytime *A* in three out of four drought-stressed saplings. Over the remaining course of the experiment, heat exposure induced continuously decreasing *A* in well-watered plants approaching those of drought-stressed plants. From the fourth day of heat onwards, no significant difference in *A* between treatments could be detected anymore (Fig. 3 a). In contrast *E* increased in response to heat in both treatments (Fig. 3 b). From the fifth day of heat onward, a slight decrease in *E* could be observed in all saplings. Significant differences between treatments of daytime *E* disappeared on the seventh day of heat, with well-watered plants reaching their minimum mean *E* of 0.4 ± 0.03 mmol m^-2^ s^-1^ on the eighth day of heat.

**Figure 3:**
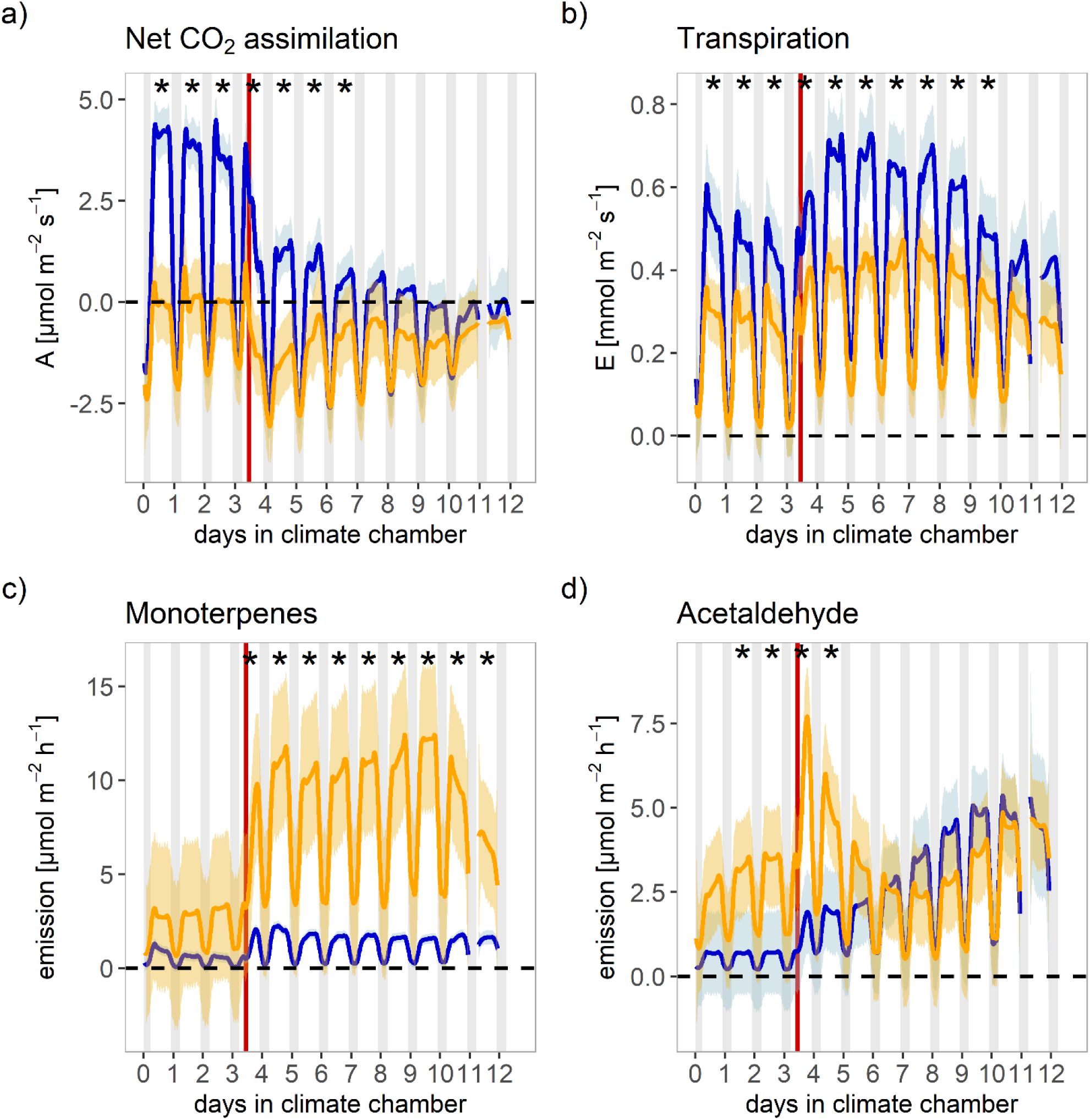
Net CO_2_ assimilation (a), transpiration (b), monoterpenes (c) and acetaldehyde emissions (d) in well-watered (blue) and drought-stressed plants (orange) in response to heat. Shaded areas indicate standard errors. Night-time is illustrated in grey and the heat ramp in red. Asterisks indicate significant differences in daily mean values between well-watered and drought-stressed plants (p ≤ 0.05). Missing values in the night of day 11 were due to technical issues.

Monoterpenes were emitted at larger rates in drought-stressed compared to well-watered plants in the pre-heat phase, even though this effect was not statistically significant (Fig. 3 c). Moreover, the synergistic effects of drought and heat further amplified that effect: upon heat exposure we observed a significant increase in monoterpene emissions of 247% and 239% in well-watered and drought-stressed plants, respectively, leading to 5-fold higher emission rates in drought-stressed plants on the first day of heat (2.1 ± 0.1 µmol m^-2^ h^-1^ in well-watered and 10.9 ± 1.4 µmol m^-2^ h^-1^ in drought-stressed plants; p ≤ 0.05). While well-watered plants maintained elevated monoterpene emission under heat, drought-stressed plants showed declining emission rates from the seventh day of heat onwards. Emissions of acetaldehyde (Fig. 3 d), isoprene and green leaf volatiles (Fig. S2) also increased immediately upon heat exposure, but followed a slightly different pattern compared to monoterpenes thereafter. Control plants and two out of four drought-stressed plants tended to increase the emission rates of these three compounds groups slightly but continuously until day 12 in the climate chamber. The other two drought-stressed plants, however, showed peak emission rates directly after the heat ramp and then continuously decreased acetaldehyde, isoprene and green leaf volatile emission (Fig. 3 d).

### Leaf terpenoid emissions

GC-MS analysis revealed 26 terpenoids emitted by needles, including 14 monoterpenes, seven additional monoterpenoids, four sesquiterpenes and one additional sesquiterpenoid. Overall, the monoterpenes β-phellandrene, limonene and β-pinene were emitted at highest rates. Emissions were dominated by monoterpenes (85% in well-watered and 93% in drought-stressed plants), followed by other monoterpenoids, sesquiterpenes and the other sesquiterpenoid. Half of the monoterpenes (β-pinene, β-phellandrene, santene, δ-3-carene, α-pinene, sabinene, p-cymene) and the sesquiterpene α-humulene were emitted in significantly higher amounts in drought-stressed plants (p ≤ 0.05), while the three monoterpenoids camphor, terpinen-4-ol and α-terpineol were emitted at higher rates in well-watered plants, though only significantly for camphor (p ≤ 0.001). For the remaining terpenoids, no significant differences were found between treatments.

Half of the compounds showed significant increases under heat compared to pre-heat (Fig. 4 a). Terpinene-4-ol and α-terpineol were emitted only in response to heat in both treatments. Total terpenoid emissions increased by 160% and 136% from the pre-heat to the second day of heat in well-watered and drought-stressed plants, respectively.

**Figure 4:**
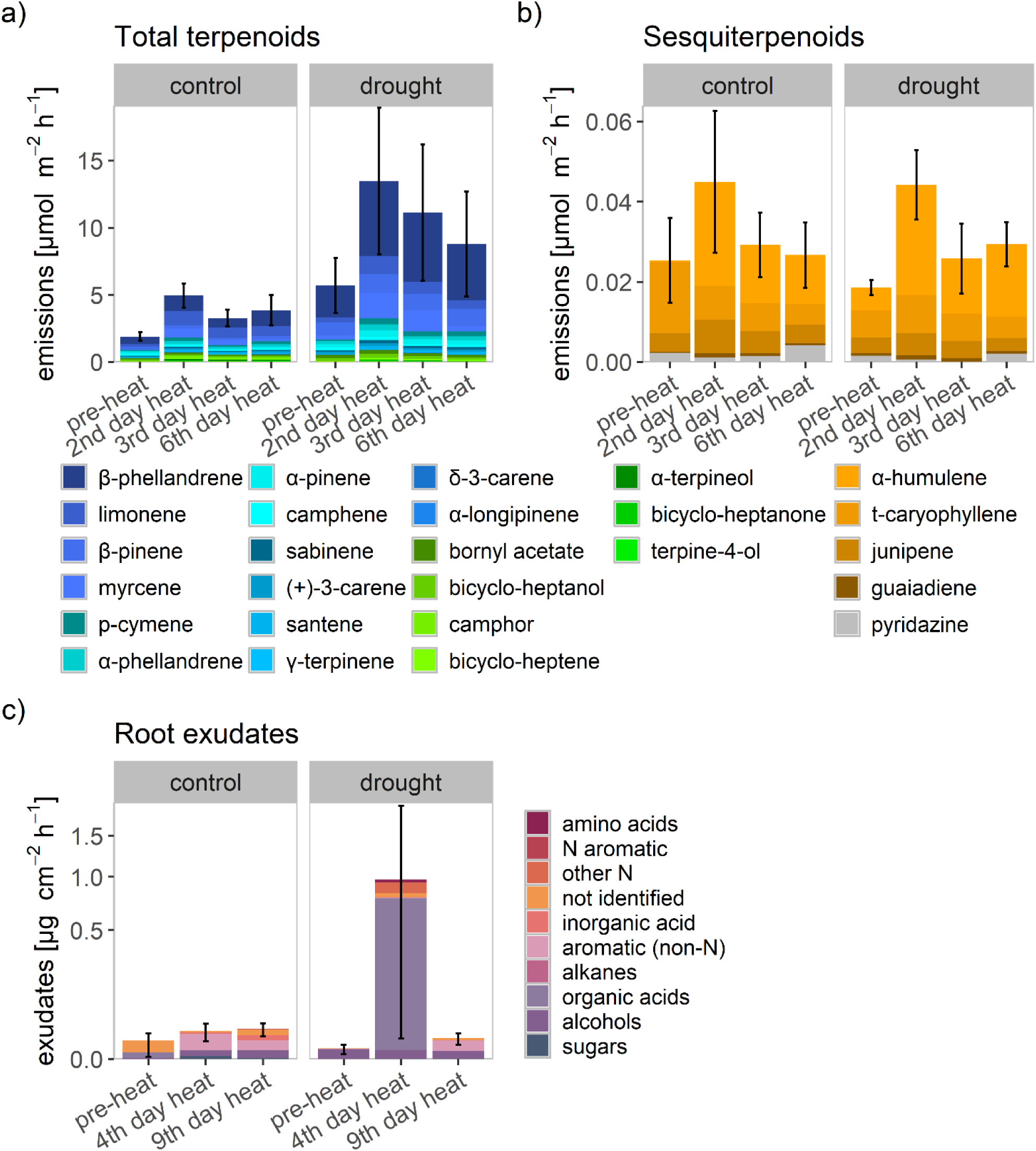
Compounds-specific total terpenoid (a) and sesquiterpenoid (b) emissions during the heat stress experiment, and root exudates (c) under heat (4^th^ day and 9^th^ day) compared to pre-heat during the drought treatment. Colours code compound groups in terpenoid emissions and root exudates. Please note the square root scales on the y-axis for root exudates.

Depending on the sampling time, drought-stressed plants showed 128% – 240% higher total terpenoid emissions than well-watered plants, which was mainly caused by an increased release of monoterpenoids, while emissions of sesquiterpenoids were comparable between the two treatments (Fig. 4 b).

### Root exudates

Overall, GC-MS analysis revealed 29 root exuded compounds (see Table S2 for complete list). In general, organic acids were exuded at highest rates, with a maximum of 10.7 µg cm^-2^ h^-1^ in drought-stressed plants in the third week of drought, and with glyoxylic acid being the most dominant compound. The least abundant compound groups were sugars and N-containing aromatics. However, the composition of exudates varied strongly between plants and sampling dates.

Heat led to an initial increase in exudation rates from roots in both treatments (Fig. 4 c). Similar to the aboveground response, drought amplified the response to heat in root exudates: the response of drought-stressed plants to heat was more pronounced, increasing exudation rates by a factor of ∼280 from pre-heat to heat, compared to a factor of ∼2 in well-watered plants. While well-watered plants maintained the same level of exudation on the ninth day of heat, the exudation rates of drought-stressed plants decreased to levels comparable to the well-watered plants before the heat application. In line with the findings from the drought treatment, during peak exudation in drought-stressed plants on the fourth day of heat, exudates were dominated by organic acids with a share of 80%. Furthermore, significantly increased sugar exudation in well-watered plants and increased exudation of aromatic compounds in both treatments could be observed the fourth day of heat compared to pre-heat values.

### Isotopic pulse labelling

The ^2^H_2_O labelling allowed to compare water uptake dynamics between the two treatments (Fig. 5 c). Δδ^2^H_2_O ratios in transpired water increased immediately after the ^2^H-labelling in both treatments, but stronger in drought-stressed plants. Significantly increased daily mean Δδ^2^H_2_O values, compared to pre-labelling values, could be observed on the following day in well-watered plants and on the second day after labelling in drought-stressed plants. In both treatments, maximum δ^2^H_2_O enrichment was reached on the fourth day after labelling (Δδ^2^H_2_O = 297.5 ± 14.4‰ for well-watered and 205.0 ± 18.5‰ for drought-stressed plants).

**Figure 5:**
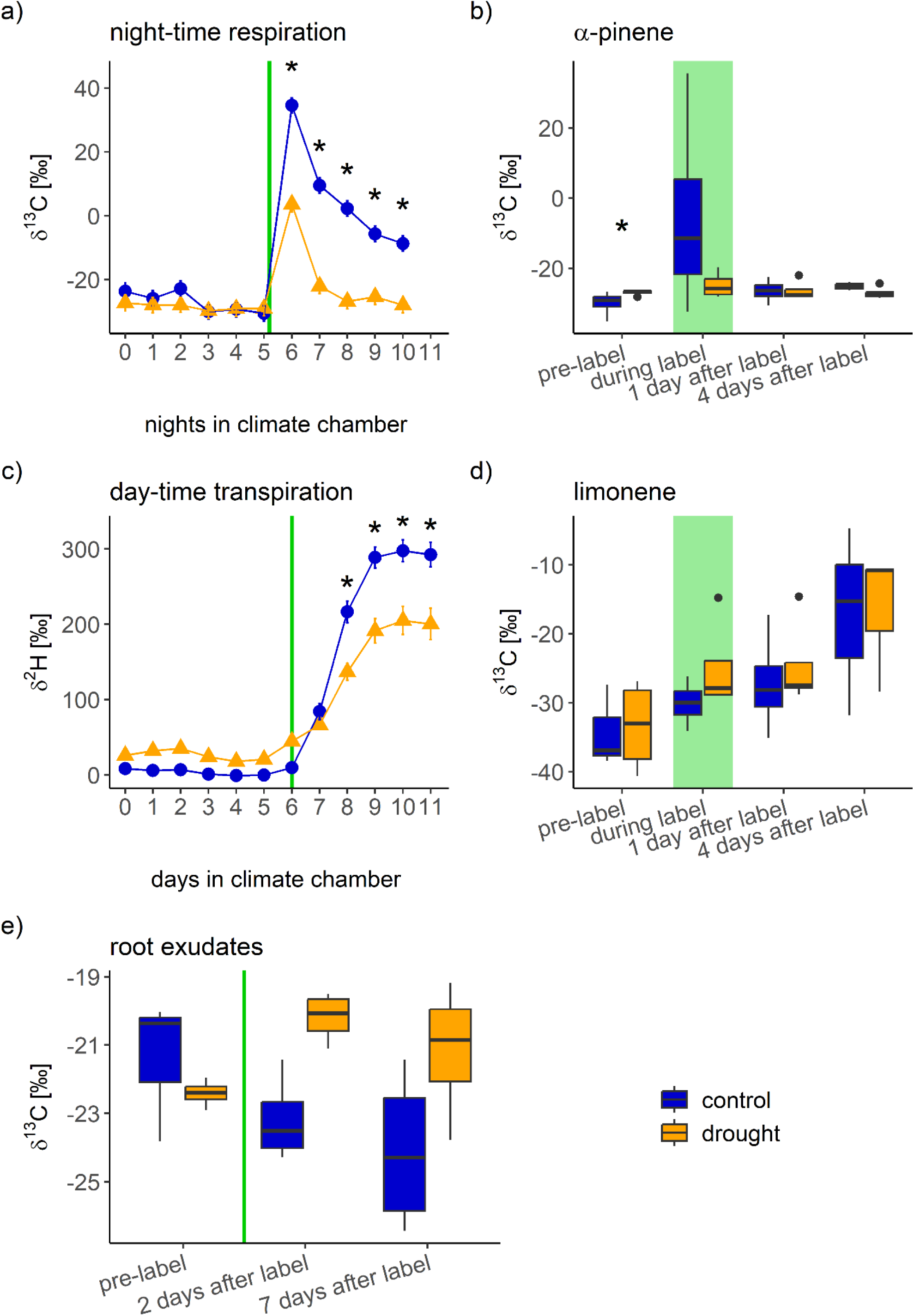
δ^13^C isotopic signatures in mean night-time respiration (a), change of δ^2^H_2_O isotopic signature (Δδ^2^H_2_O) in mean day-time transpiration as compared to pre-label values of well-watered plants (c), δ^13^C isotopic signature in α-pinene (b) and limonene (d) emissions, and root exudates (e). The green bars mark the time of labelling. Asterisks indicate significant differences between treatments.

The natural abundance δ^13^CO_2_ of night-time respiration was constant and not significantly different between the two treatments (-29.3 ± 2.4‰ for well-watered, -29.2 ± 2.4‰ for drought-stressed plants; Fig. 5 a). In the night following the label event, δ^13^CO_2_ in respiration significantly increased in both treatments, but much less in drought-stressed plants (+34.6 ± 2.4‰ for well-watered and +3.6 ± 2.4‰ for drought-stressed plants). In drought-stressed plants, δ^13^CO_2_ in respiration declined to initial values already in the second night after labelling, while in well-watered plants, values also decreased, but remained elevated compared to pre-labelling and compared to drought-stressed plants.

The ^13^CO_2_-labelling led only to a slight increase in δ^13^C ratios in VOCs, but allowed to categorise the emitted compounds into three groups: The first group showed the highest label incorporation during the ^13^CO_2_ application and successively decreased thereafter. This pattern could be observed in α-pinene (Fig. 5 b), sabinene, myrcene, α-terpineol and p-cymene. The second group showed steadily increasing δ^13^C values reaching the highest levels four days after the labelling event and was represented by limonene (Fig. 5 d) and junipene. Bornyl acetate, β-pinene and camphene showed constant, non-elevated δ^13^C ratios in VOC emissions, representing the third group.

δ^13^C ratios in root exudates varied between -20‰ and -24‰ prior to ^13^CO_2_ application (Fig. 5 e). While values in drought-stressed plants tended to increase reaching δ^13^C ratios up to -19‰, δ^13^C tended to decrease in exudates from well-watered plants after the ^13^CO_2_ exposure with a maximum of around -21‰,. The decrease in δ^13^C in the exudates of well-watered plants was most likely due to the exposure to the cuvette air, which is supplied by anthropogenic CO_2_, which is more depleted in ^13^C compared to the air outside (δ^13^CO_2_ = -35.26 ± 0.03‰ in the cuvette air).

## Discussion

Our study demonstrates the severe effects of heat, drought and their interaction on the physiology of Norway spruce saplings. In particular, the interaction of heat and drought induced a synergistic effect on leaf VOC emission and root exudate rates. This means that the C loss under interacting heat and drought exceeded the sum of C losses from single stressors, which might lead to unexpected releases under compound droughts. Isotopic labelling indicated that heat-stressed plants invested higher proportions of freshly assimilated C into VOC synthesis, while plants exposed to combined drought and heat rather relied on stored VOCs. Inversely, root exudates were supplied with freshly assimilated C in heat- and drought-stressed plants, whereas heat alone did not lead to *de novo* C investment into root exudates. This clearly demonstrates the high responsiveness to environmental stressors and non-linear interaction effects, which will be discussed in the following.

### Synergistic effect of heat and drought on C losses despite plummeting photosynthetic activity

Drought exposure significantly reduced assimilation and transpiration rates of Norway spruce saplings with assimilation experiencing greater relative reductions compared to transpiration. The enrichment of δ^13^CO_2_ upon labelling in the respiration of drought-stressed plants indicates that they were still photosynthetically active despite negative net *A* rates. However, the stronger, long-lasting effect of the ^13^CO_2_ pulse labelling on the respiration of well-watered plants demonstrate their higher capacity in terms of C uptake, which also holds for water uptake. Even though plants received the same amount of labelled water in both treatments, well-watered plants showed generally higher label strengths indicating a higher capacity for uptake and transport of water.

Our results show that Norway spruce increases the diffusive resistance of CO_2_ through the closure of stomata (Deslauriers *et al*., 2014; Daber *et al*., 2025). Heat also affects the photosynthetic activity by increasing photorespiration and deactivating Rubisco (Rennenberg *et al*., 2006; Filella *et al*., 2007) resulting in decreasing *A* under heat in Norway spruce as reported elsewhere (Deslauriers *et al*., 2014; Meischner *et al*., 2024). In well-watered plants, transpiration is important for evaporative cooling, strongly mitigating the effect of heat. However, in drought-stressed plants, transpiration in reduced, enhancing negative consequences of elevated air temperature (Eamus *et al*., 2008), which might explain amplified responses in VOC emissions. VOC emissions as well as root exudates are known to increase stress tolerance. Our results clearly show the stimulation of these processes under drought and heat, despite negative *A* values. While heat and drought caused similar increases in VOC emissions compared to non-stress conditions, exudation rates from roots reacted stronger to heat than to drought, reflecting the high sensitivity of Norway spruce to heat (Kunert, 2020; Jamnická *et al*., 2024). Interestingly, we observed a synergistic effect of heat and drought on VOC emission and exudation rates from roots, i.e. release rates under combined stress were larger than the sum of the release rates under single stresses. Especially exudation rates from roots increased drastically under combined heat and drought, in spite of negative *A* rates. However, the increased investment in VOC emissions and root exudates could not be sustained by the plants. After the fourth day of heat, VOC emission and exudation rates decreased again, even though they were still elevated, which might be due to C depletion, especially since photosynthesis continued to decrease. Many studies have already stressed the complexity of multiple stress situations in forest trees (Callaway and Walker, 1997; Brooker, 2006; Doblas-Miranda *et al*., 2017) and it has been suggested that only few tree species are tolerant to two or more co-occurring stressors (Niinemets, 2010). Possibilities for stress interaction in natural environments are tremendous (Côté *et al*., 2016) and our data illustrate the detrimental effect of increasingly frequent compound events on Norway spruce saplings.

### Increased VOC emissions under heat and drought in Norway spruce

Leaf VOC emissions in Norway spruce saplings increased under drought. This contradicts the findings of Daber *et al*. (2025), who found decreased terpenoid emissions in Norway spruce under drought. Saplings in the experiment of Daber *et al*. (2025) were younger (2 years) and potentially had smaller storage pools, which might explain the contrasting results; especially since there were no immediate increases in VOC-δ^13^C in drought-stressed plants and *A* was strongly reduced in both studies. Increased monoterpenoid emissions in drought- and heat-stressed plants, despite reduced photosynthetic rates and limited label incorporation, suggest a high importance of VOC storage pools under multiple stressors. VOC storage pools are known to be especially large in Norway spruce (Fischbach *et al*., 2000). VOC emissions are known to increase under drought initially (Loreto and Schnitzler, 2010) and may protect already stressed plants against herbivory e.g., bark beetle infestation (Lehmanski *et al*., 2023). In contrast to monoterpenoids, sesquiterpenoid emission rates were not affected by drought in our study, suggesting that the mevalonic acid (MVA) pathway (Dudareva *et al*., 2013) was not affected by drought, while the methylerythritol phosphate (MEP) pathway synthesising monoterpenoids (Dudareva *et al*., 2013) was upregulated. Interestingly, the only three compounds, which were emitted in higher rates in well-watered compared to drought-stressed plants were oxygenated monoterpenoids (camphor, terpinen-4-ol and α-terpineol). Additionally, the emission rates of those compounds increased under heat. These patterns might indicate an interrelation of oxygenated monoterpenoid emissions with transpiration rates as suggested earlier by Duan *et al*. (2020), who detected oxygenated terpenoids in the xylem sap being transported to the leaves.

We observed a clear increase of monoterpene emissions under heat. The temperature dependence of VOC emissions is well known (Guenther *et al*., 1993) and especially high in Norway spruce compared to other species (Filella *et al*., 2007). Norway spruce has shown an exponential increase of monoterpene emissions with temperature (Grabmer *et al*., 2006; Filella *et al*., 2007; Meischner *et al*., 2024). This can be explained by an increased monoterpene synthase activity, which has its optimum temperature at 40°C in Norway spruce, and higher volatility (Fischbach *et al*., 2000). This is backed up by C-label incorporation in VOCs of well-watered plants under heat. Immediate label incorporation into VOCs could be detected for α-pinene, sabinene and myrcene. Even though their enrichment in δ^13^C was not significant, it indicates that these compounds were partially synthesised *de novo*. Increased VOC emission rates under heat serve to induce signalling cascades to stabilise membranes and to quench reactive oxygen species (Sharkey *et al*., 2008; Loreto and Schnitzler, 2010).

In contrast to the amplification of VOC emissions we observed, a buffering effect of heat on VOC emissions in the presence of multiple stressors (heat and herbivory) has been observed and reasoned by reduced photosynthesis rates by Meischner *et al*. (2024). Similarly, a simulation of monoterpene emissions by Rennenberg *et al*. (2006) in oak and beech under heat with and without drought showed reduced emission rates under combined stress compared to heat alone. However, oak and beech do not have large terpenoid storage pools such as spruce (Duan *et al*., 2020). Underlining this, we detected no label incorporation in bornyl acetate, β-pinene and camphene suggesting their emissions from storage pools. Besides, there is a group of VOCs that showed the peak of label incorporation not immediately upon labelling but delayed. This applied to limonene and junipene, and suggests short-term storage of these compounds before emission or the synthesis with assimilates from previous days. That terpenoid emission rates finally decreased under combined heat and drought could may be due to C starvation, which is supported by negative *A* already in the days before emission rates started to decrease.

If compound droughts (Werner *et al*., 2025) result in a synergistic effect on VOC emissions as observed in our study also in natural environments, implications for atmospheric effects may be larger than foreseen so far. Negative net assimilation rates accompanied by increased emission rates could make forests to significant C sources to the atmosphere during compound events, adding to the consequences of VOC emissions on atmospheric chemistry mentioned earlier. Next to releases to the atmosphere, stress situations also cause C losses from temperate forest trees to the soil, for example in the form of root exudates.

### Root exudates under heat and drought in Norway spruce

So far, no consensus on the effect of drought on root exudates has been reached with increases (Meier *et al*., 2020; Leuschner *et al*., 2022) as well as decreases (Dannenmann *et al*., 2009; Li *et al*., 2025) being reported in literature. Gargallo-Garriga *et al*. (2018) suggested that exudation is increased under mild drought to facilitate soil penetration, while exudation is reduced under prolonged or severe droughts to save resources. We observed highest exudation rates in drought-stressed plants under heat, which indicates an amplifying effect of heat on exudation from roots under drought. However, the duration of drought could also play a role considering that exudation rates were highest at the onset of drought, which coincided with the natural heat wave, and decreased thereafter. Nonetheless, heat caused an increase in exudation rates, also in well-watered plants, and amplified the effect of drought, which was similarly observed under controlled heat. An increase in exudation rates under elevated temperature has already been observed in previous studies (Leuschner *et al*., 2022; Heinzle *et al*., 2023). However, this increase in exudation rates from roots under heat seems to be a short-term adaptation, which is not maintained under continued heat (Leuschner *et al*., 2022; Liu *et al*., 2025). It has been postulated that root exudates could be a short-term mechanism to dispose of surplus C assimilated in a warmer environment which cannot be used instantly due to a lack of water or nutrients (Prescott *et al*., 2020). However, given the large increase in exudation rates under combined heat and drought, while net C-uptake was negative, it seems unlikely that exudates were driven by surplus C. Our data rather suggest exudation from roots to be an active stress compensation response, which has also been shown for VOC emissions (Loreto and Schnitzler, 2010). Moreover, increased organic acid exudation has been reported under P (Jones *et al*., 2009; Vives-Peris *et al*., 2020) and N (Tückmantel *et al*., 2017; Vives-Peris *et al*., 2020) deficiency. The observed increase in organic acid exudation under heat, partially combined with drought, might reflect a compensation for impeded nutrient uptake conditions under heat and drought. We also observed an increased exudation of sugars and aromatic compounds under heat, which have been identified to facilitate Fe uptake and might therefore serve a similar purpose as organic acids (Vranova *et al*., 2013; Vives-Peris *et al*., 2020). We are still lacking a sound knowledge on the function of specific root exudates (Ritter *et al*., 2025); however, our results suggest that the exudation of sugars and aromatic compounds may aid to increase heat tolerance.

Drought-stressed plants tended to dedicate more of *de novo* synthesised compounds to root exudates compared to well-watered plants. Even though effects were not statistically significant, they are supported by studies showing an increased investment in belowground organs upon drought (Oppenheimer-Shaanan *et al*., 2022). Furthermore, Brunn *et al*. (2025) observed that even under reduced assimilation under drought, the downward C-flux within the tree stayed the same, indicating a higher share of freshly assimilated C is allocated to root exudates. Well-watered plants did not show *de novo* assimilated C investment into root exudates, which suggests that under heat with sufficient water supply, newly assimilated C is prioritised to other sinks than root exudates, e.g. leaf VOC emissions.

### Conclusion

Our results shed light on how resources are partitioned to above and belowground processes in response to heat and drought in Norway spruce saplings. Specifically, we observed that drought increased VOC emissions, while exudation rates from roots were reduced. Heat had a similar effect on VOC emissions as drought, but also increased exudation from root. Finally, we could show clear synergistic effects of heat and drought, resulting in amplified responses of both, VOC emissions and exudation, even though photosynthetic assimilation was strongly reduced with negative net carbon gain. This is highly relevant given the projected increase of compound droughts under future climate conditions (Werner *et al*., 2025).

Our experiment enabled insights to the simultaneous above- and belowground stress response of Norway spruce under multiple stressors, which is widely unstudied in forest trees. Linking processes from the above- and belowground responses may yield a more holistic understanding of ecophysiological functioning in response to compound droughts in the future.

## Supplementary data

The following supplementary data are available at JXB online:

Table S1: Complete list of leaf-emitted terpenoids.

Table S2: Complete list of root exudates.

Fig. S1: Net assimilation and transpiration rates during the drought treatment.

Fig. S2: Isoprene and green leaf volatile emissions during the heat stress experiment.

## Acknowledgements

This study was conducted in the framework of the Research Unit 5315 “Forest Floor: Functioning, Dynamics and Vulnerability in a Changing World” funded by the DFG (WE2681/13-1) and the Collaborative Research Centre 1537 “ECOSENSE Multiscale quantification of spatio-temporal dynamics of ecosystem processes by smart autonomous sensor networks” (459819582). We thank A. Paul, R. Schneider and E. Schottmüller for lab assistance, analyses and gardening, and the students L. Wolz and H. Vogt for sampling.

## Author contributions

MW, MM, CS, SD, SH and CW designed the study. MM, CS, SD and MW set up the experiment. MW, CS and SD organised and conducted measurements. MW conducted samplings and lab analyses with assistance of JK. MW analysed the data with major contributions of MM and guidance of SH and CW and drafted the manuscript. All authors reviewed the manuscript and contributed substantially.

## Data availability

The data supporting this study is available from the corresponding author upon reasonable request.

## Abbreviations

A: net assimilation rate
E: transpiration rate
Fig.: figure
VOC: volatile organic compound
VWC: soil volumetric water content
Ψ_leaf_: leaf water potential

## References

Arend M, Link RM, Patthey R, Hoch G, Schuldt B, Kahmen A. 2021. Rapid hydraulic collapse as cause of drought-induced mortality in conifers. Proceedings of the National Academy of Sciences of the United States of America 118.

Atkinson R, Arey J. 2003. Gas-phase tropospheric chemistry of biogenic volatile organic compounds: a review. Atmospheric Environment 37, 197–219.

Bevacqua E, Zappa G, Lehner F, Zscheischler J. 2022. Precipitation trends determine future occurrences of compound hot–dry events. Nature Climate Change 12, 350–355.

Bonn B, Magh R-K, Rombach J, Kreuzwieser J. 2019. Biogenic isoprenoid emissions under drought stress: different responses for isoprene and terpenes. Biogeosciences 16, 4627–4645.

Bourtsoukidis E, Pozzer A, Williams J et al. 2024. High temperature sensitivity of monoterpene emissions from global vegetation. Communications Earth & Environment 5, 23.

Brunn M, Hafner BD, Zwetsloot MJ et al. 2022. Carbon allocation to root exudates is maintained in mature temperate tree species under drought. New phytologist 235, 965–977.

Brunn M, Mueller CW, Chari NR et al. 2025. Tree carbon allocation to root exudates: implications for carbon budgets, soil sequestration and drought response. Tree Physiology 45.

Caemmerer S von, Farquhar GD. 1981. Some relationships between the biochemistry of photosynthesis and the gas exchange of leaves. Planta 153, 376–387.

Côté IM, Darling ES, Brown CJ. 2016. Interactions among ecosystem stressors and their importance in conservation. Proceedings. Biological sciences 283.

Daber LE, Nolte P, Kreuzwieser J, Meischner M, Williams J, Werner C. 2025. Position-specific isotope labelling gives new insights into chiral monoterpene synthesis of Norway spruce (Picea abies L.). Environmental and Experimental Botany 238, 106238.

Dannenmann M, Simon J, Gasche R et al. 2009. Tree girdling provides insight on the role of labile carbon in nitrogen partitioning between soil microorganisms and adult European beech. Soil Biology and Biochemistry 41, 1622–1631.

Deslauriers A, Beaulieu M, Balducci L, Giovannelli A, Gagnon MJ, Rossi S. 2014. Impact of warming and drought on carbon balance related to wood formation in black spruce. Annals of Botany 114, 335–345.

Duan Q, Bonn B, Kreuzwieser J. 2020. Terpenoids are transported in the xylem sap of Norway spruce. Plant, Cell & Environment 43, 1766–1778.

Dubbert M, Cuntz M, Piayda A, Werner C. 2014. Oxygen isotope signatures of transpired water vapor: the role of isotopic non-steady-state transpiration under natural conditions. The New phytologist 203, 1242–1252.

Dudareva N, Klempien A, Muhlemann JK, Kaplan I. 2013. Biosynthesis, function and metabolic engineering of plant volatile organic compounds. The New phytologist 198, 16–32.

Eamus D, Taylor DT, Macinnis-Ng CMO, Shanahan S, Silva L de. 2008. Comparing model predictions and experimental data for the response of stomatal conductance and guard cell turgor to manipulations of cuticular conductance, leaf-to-air vapour pressure difference and temperature: feedback mechanisms are able to account for all observations. Plant, Cell & Environment 31, 269–277.

Fasbender L, Yáñez-Serrano AM, Kreuzwieser J, Dubbert D, Werner C. 2018. Real-time carbon allocation into biogenic volatile organic compounds (BVOCs) and respiratory carbon dioxide (CO2) traced by PTR-TOF-MS, 13CO2 laser spectroscopy and 13C-pyruvate labelling. PloS one 13, e0204398.

Filella I, Wilkinson MJ, Llusià J, Hewitt CN, Peñuelas J. 2007. Volatile organic compounds emissions in Norway spruce (Picea abies) in response to temperature changes. Physiologia plantarum 130, 58–66.

Fischbach RJ, Zimmer I, Steinbrecher R, Pfichner A, Schnitzler JP. 2000. Monoterpene synthase activities in leaves of Picea abies (L.) Karst. and Quercus ilex L. Phytochemistry 54, 257–265.

Gargallo-Garriga A, Preece C, Sardans J, Oravec M, Urban O, Peñuelas J. 2018. Root exudate metabolomes change under drought and show limited capacity for recovery. Scientific Reports 8, 12696.

Gazol A, Camarero JJ. 2022. Compound climate events increase tree drought mortality across European forests. The Science of the total environment 816, 151604.

Grabmer W, Kreuzwieser J, Wisthaler A et al. 2006. VOC emissions from Norway spruce (Picea abies L. [Karst]) twigs in the field—Results of a dynamic enclosure study. Atmospheric Environment 40, 128–137.

Guenther AB, Zimmerman PR, Harley PC, Monson RK, Fall R. 1993. Isoprene and monoterpene emission rate variability: Model evaluations and sensitivity analyses. Journal of Geophysical Research: Atmospheres 98, 12609–12617.

Haberstroh S, Kreuzwieser J, Boeddeker H et al. 2019. Natural Carbon Isotope Composition Distinguishes Compound Groups of Biogenic Volatile Organic Compounds (BVOC) in Two Mediterranean Woody Species. Frontiers in Forests and Global Change 2, 463563.

Haberstroh S, Kreuzwieser J, Lobo-do-Vale R, Caldeira MC, Dubbert M, Werner C. 2018. Terpenoid Emissions of Two Mediterranean Woody Species in Response to Drought Stress. Frontiers in plant science 9, 1071.

Hakola H, Tarvainen V, Praplan AP et al. 2017. Terpenoid and carbonyl emissions from Norway spruce in Finland during the growing season. Atmospheric Chemistry and Physics 17, 3357–3370.

Heinzle J, Liu X, Tian Y et al. 2023. Increase in fine root biomass enhances root exudation by long-term soil warming in a temperate forest. Frontiers in Forests and Global Change 6, 1152142.

Hesse BD, Hikino K, Gebhardt T et al. 2024. Acclimation of mature spruce and beech to five years of repeated summer drought - The role of stomatal conductance and leaf area adjustment for water use. The Science of the total environment 951, 175805.

Holopainen JK. 2004. Multiple functions of inducible plant volatiles. Trends in plant science 9, 529–533.

Jamnická G, Húdoková H, Fleischer P, Ježík M. 2024. Soil drought stress and high-temperature effects on photosystem II in different juvenile spruce provenances. Central European Forestry Journal 70, 95–106.

Jones DL, Nguyen C, Finlay RD. 2009. Carbon flow in the rhizosphere: carbon trading at the soil–root interface. Plant and Soil 321, 5–33.

Jordan A, Haidacher S, Hanel G et al. 2009. A high resolution and high sensitivity proton-transfer-reaction time-of-flight mass spectrometer (PTR-TOF-MS). International Journal of Mass Spectrometry 286, 122–128.

Karpov A, Pirtskhalava-Karpova N, Singh VV et al. 2026. Response of mature Norway spruce to experimental thermal and drought stress in relation to Ips typographus attack: Crown temperatures and sap flow. Forest Ecology and Management 600, 123290.

Knoke T, Gosling E, Thom D, Chreptun C, Rammig A, Seidl R. 2021. Economic losses from natural disturbances in Norway spruce forests – A quantification using Monte-Carlo simulations. Ecological Economics 185, 107046.

Kolář T, Čermák P, Trnka M, Žid T, Rybníček M. 2017. Temporal changes in the climate sensitivity of Norway spruce and European beech along an elevation gradient in Central Europe. Agricultural and Forest Meteorology 239, 24–33.

Kreuzwieser J, Hauberg J, Howell KA et al. 2009. Differential response of gray poplar leaves and roots underpins stress adaptation during hypoxia. Plant physiology 149, 461–473.

Kreuzwieser J, Meischner M, Grün M, Yáñez-Serrano AM, Fasbender L, Werner C. 2021. Drought affects carbon partitioning into volatile organic compound biosynthesis in Scots pine needles. The New phytologist 232, 1930–1943.

Kunert N. 2020. Preliminary indications for diverging heat and drought sensitivities in Norway spruce and Scots pine in Central Europe. iForest - Biogeosciences and Forestry 13, 89–91.

Kunert N, Hajek P, Hietz P, Morris H, Rosner S, Tholen D. 2022. Summer temperatures reach the thermal tolerance threshold of photosynthetic decline in temperate conifers. Plant biology (Stuttgart, Germany) 24, 1254–1261.

Lehmanski LMA, Kandasamy D, Andersson MN et al. 2023. Addressing a century-old hypothesis - do pioneer beetles of Ips typographus use volatile cues to find suitable host trees? The New phytologist 238, 1762–1770.

Leuschner C, Tückmantel T, Meier IC. 2022. Temperature effects on root exudation in mature beech (Fagus sylvatica L.) forests along an elevational gradient. Plant and Soil 481, 147–163.

Li W, Zhang H, Pu Y, Li F. 2025. Effects of drought on carbon-nitrogen dynamics in root exudates of Qinghai spruce. Tree Physiology 45.

Liu X, Heinzle J, Tian Y et al. 2025. Primary metabolites in root exudates are not affected by long-term soil warming in a temperate forest. Functional Ecology.

Loreto F, Schnitzler J-P. 2010. Abiotic stresses and induced BVOCs. Trends in plant science 15, 154–166.

Maurer D, Malique F, Alfarraj S et al. 2021. Interactive regulation of root exudation and rhizosphere denitrification by plant metabolite content and soil properties. Plant and Soil 467, 107–127.

Meier IC, Tückmantel T, Heitkötter J et al. 2020. Root exudation of mature beech forests across a nutrient availability gradient: the role of root morphology and fungal activity. New phytologist 226, 583–594.

Meischner M, Dumberger S, Daber LE et al. 2024. Jasmonic acid and heat stress induce high volatile organic compound emissions in Picea abies from needles, but not from roots. Tree Physiology.

Meischner M, Haberstroh S, Kreuzwieser J et al. 2025. Localized Response of De Novo Terpenoid Emissions Through the Jasmonate Signaling Cascade in Two Main European Tree Species. Physiologia plantarum 177, e70432.

Niinemets Ü. 2010. Responses of forest trees to single and multiple environmental stresses from seedlings to mature plants: Past stress history, stress interactions, tolerance and acclimation. Forest Ecology and Management 260, 1623–1639.

Oberhuber W, Hammerle A, Kofler W. 2015. Tree water status and growth of saplings and mature Norway spruce (Picea abies) at a dry distribution limit. Frontiers in plant science 6, 703.

Oppenheimer-Shaanan Y, Jakoby G, Starr ML et al. 2022. A dynamic rhizosphere interplay between tree roots and soil bacteria under drought stress. eLife 11.

Ormeño E, Mévy JP, Vila B et al. 2007. Water deficit stress induces different monoterpene and sesquiterpene emission changes in Mediterranean species. Relationship between terpene emissions and plant water potential. Chemosphere 67, 276–284.

Peñuelas J, LLusià J, Asensio D, Munné-Bosch S. 2005. Linking isoprene with plant thermotolerance, antioxidants and monoterpene emissions. Plant, Cell & Environment 28, 278–286.

Phillips RP, Erlitz Y, Bier R, Bernhardt ES. 2008. New approach for capturing soluble root exudates in forest soils. Functional Ecology 22, 990–999.

Prescott CE, Grayston SJ, Helmisaari H-S et al. 2020. Surplus Carbon Drives Allocation and Plant-Soil Interactions. Trends in Ecology & Evolution 35, 1110–1118.

Rennenberg H, Loreto F, Polle A et al. 2006. Physiological responses of forest trees to heat and drought. Plant biology (Stuttgart, Germany) 8, 556–571.

Ritter A, Becker PJ, Möller K et al. 2025. Targeting the untargeted: Uncovering the chemical complexity of root exudates. Plant and Soil, 1–16.

Rohrbacher F, St-Arnaud M. 2016. Root Exudation: The Ecological Driver of Hydrocarbon Rhizoremediation. Agronomy 6, 19.

Sharkey TD, Wiberley AE, Donohue AR. 2008. Isoprene emission from plants: why and how. Annals of Botany 101, 5–18.

Tückmantel T, Leuschner C, Preusser S et al. 2017. Root exudation patterns in a beech forest: Dependence on soil depth, root morphology, and environment. Soil Biology and Biochemistry 107, 188–197.

Ulrich DEM, Grossiord C. 2023. Faster drought recovery in anisohydric beech compared with isohydric spruce. Tree physiology 43, 517–521.

Vives-Peris V, Ollas C de, Gómez-Cadenas A, Pérez-Clemente RM. 2020. Root exudates: from plant to rhizosphere and beyond. Plant cell reports 39, 3–17.

Vranova V, Rejsek K, Skene KR, Janous D, Formanek P. 2013. Methods of collection of plant root exudates in relation to plant metabolism and purpose: A review. Journal of Plant Nutrition and Soil Science 176, 175–199.

Wannenmacher M, Haberstroh S, Kreuzwieser J et al. 2025. Spatio-temporal plasticity of root exudation in three temperate tree species: effects of season, site and soil characteristics.

Werner C, Bahn M, Grams TEE et al. 2025. Impact of emerging compound droughts on forests: A water supply and demand perspective. Plant biology (Stuttgart, Germany).

Werner C, Fasbender L, Romek KM, Yáñez-Serrano AM, Kreuzwieser J. 2020. Heat Waves Change Plant Carbon Allocation Among Primary and Secondary Metabolism Altering CO2 Assimilation, Respiration, and VOC Emissions. Frontiers in Plant Science 11, 1242.

Werner C, Meredith LK, Ladd SN et al. 2021. Ecosystem fluxes during drought and recovery in an experimental forest. Science (New York, N.Y.) 374, 1514–1518.

Werner C, Wegener F, Unger S, Nogués S, Priault P. 2009. Short-term dynamics of isotopic composition of leaf-respired CO2 upon darkening: measurements and implications. Rapid communications in mass spectrometry RCM 23, 2428–2438.

Wickham H. 2016. ggplot2. Cham: Springer International Publishing.

Wu C, Pullinen I, Andres S et al. 2015. Impacts of soil moisture on de novo monoterpene emissions from European beech, Holm oak, Scots pine, and Norway spruce. Biogeosciences 12, 177–191.

Yáñez-Serrano AM, Filella I, LLusià J et al. 2021. GLOVOCS - Master compound assignment guide for proton transfer reaction mass spectrometry users. Atmospheric Environment 244, 117929.

